# High-contrast multifocus microscopy with a single camera and z-splitter prism

**DOI:** 10.1101/2020.08.04.236661

**Authors:** Sheng Xiao, Howard Gritton, Hua-an Tseng, Dana Zemel, Xue Han, Jerome Mertz

## Abstract

We present a multifocus imaging strategy based on the use of a simple z-splitter prism that can be assembled from off-the-shelf components. Our technique enables a widefield image stack to be distributed onto a single camera and recorded simultaneously. We exploit the volumetric nature of our image acquisition by further introducing a novel extended-volume 3D deconvolution strategy to suppress far-out-of-focus fluorescence background to significantly improve the contrast of our recorded images, conferring to our system a capacity for quasi optical sectioning. By swapping in different z-splitter configurations, we can prioritize high speed or large 3D field-of-view imaging depending on the application of interest. Moreover, our system can be readily applied to a variety of imaging modalities in addition to fluorescence, such as phase-contrast and darkfield imaging. Because of its simplicity, versatility, and performance, we believe our system will be a useful tool for general biological or biomedical imaging applications.

## 1 Introduction

Optical microscopy has been an indispensable tool for studying 3D complex biological systems^1^. By far, the most common technique to record images with a microscope is with a digital camera, which is cost effective, light efficient, low noise, simple to use and involves no moving parts. Moreover, with current sCMOS technology, cameras offer unparalleled sampling speeds (several kHz) and pixel counts (several megapixels). However, standard camera-based optical microscopes provide sharp, in-focus imaging only of a single 2D plane, with out-of-focus sample structure becoming increasingly blurred with in-focus sharpness because of the inherent trade-off between spatial resolution and depth-of-field^2^. For thick samples, such out-of-focus blurring produces background that undermines the contrast and signal-to-noise ratio (SNR) of the in-focus structure.

Several strategies have been developed to enable camera-based microscopes to produce sharp images over extended depth ranges^3^. The simplest, of course, is to acquire multiple images while translating the sample and/or camera sensor. This is also the slowest and most unwieldy. Considering that biological processes often occur on the millisecond time scale, faster image-stack acquisition is generally desired, which can be obtained, for example, by remote focusing with a fast z-scanner^4,5^. Image stacks can also be obtained in light-sheet geometries^6–8^ with the benefit of reduced background generation. These strategies involve multiple sequential image acquisitions, and thus their volumetric imaging rate is inherently reduced compared to the native frame rate of the camera.

To obtain instantaneous (single frame) imaging over an extended volume, different strategies can be considered. Quasi-3D, or extended-depth-of-field (EDOF), imaging can be obtained through single 2D projections by means of ultrafast focus scans^9–12^ or point spread function (PSF) engineering^13–15^, albeit with loss of axial resolution. Full 3D information can be retrieved by combining computational techniques with depth-dependent PSFs^16–19^, however this generally requires sample sparsity or other *a priori* information. For thick or densely labeled samples, such 2D projection techniques are undermined by loss of image contrast, owing to the broadening of PSFs and spatial superposition of multiple images from different focal planes. In these cases, out-of-focus background suppression performed physically^20,21^ or numerically^22,23^ is beneficial.

Alternatively, instantaneous volumetric information can be captured by simultaneously recording distinct images from multiple focal planes. Such a multifocus strategy avoids spatially superposing distinct images and has the advantage of preserving sharp in-focus PSFs at each focal plane, making it better suited for imaging dense volumetric samples. Simultaneous multifocus imaging can be performed with multiple cameras^24–26^, or, more practically, by image splitting onto a single camera, taking advantage of the enormous pixel numbers afforded by modern sCMOS sensors. In the latter case, an optical device is required to convert axially distributed focal planes within the sample to transversely distributed focal planes at the camera sensor. To date, examples of such devices include specially fabricated beam splitters (using two cameras^27,28^) or diffractive optical elements (DOEs)^29–32^. These usually involve significant design and manufacturing challenges. Moreover, their applications have been mostly limited to super-resolution imaging and single particle tracking in thin samples over relatively small 3D volumes.

Here, we describe a simple and flexible camera-based microscopy approach for high-speed, large field-of-view multifocus imaging of live cells and organisms. Our technique makes use of a low-cost and light efficient multiplane-imaging prism, called a z-splitter, that splits the detection light path into a sequence of multiple paths directed onto a single camera, such that each detection path conjugates a different focal plane. In this manner, an axially distributed sequence of focal planes in the sample is simultaneously imaged in a single camera frame. We provide three different z-splitter geometries, generalized from the same basic design and made entirely from off-the-shelf components, that can be readily swapped in and out of our microscope depending on whether we wish to prioritize speed (> 3 × the camera full-frame rate) or large imaging volumes (up to 9 focal planes spanning a 1.1 × 1.1 × 0.7 mm^3^ volume).

Our technique is based on standard widefield microscopy and, as such, does not inherently provide optical sectioning^1^. Nevertheless, we exploit the fact that our system images multiple focal planes simultaneously, enabling quasi optical-sectioning by numerical suppression of out-of-focus background^33^. To this end, a large portion of our work is devoted to the development and implementation of a novel 3D deconvolution strategy that explicitly extrapolates the reconstruction of the fluorescent sample beyond the volume actually spanned by the imaged focal planes. We show that such extended-volume 3D (EV-3D) deconvolution significantly improves background suppression, leading to high image contrast and signal-to-noise-ratio (SNR) even in thick samples such as densely labeled mouse brain. By making use of a warm-start strategy, fast EV-3D deconvolution can be performed at speeds of 0.4 s per volume.

Finally, we demonstrate that our z-splitting technique is easily compatible with a variety of imaging modalities, including fluorescence, phase-contrast and darkfield imaging, making it a highly versatile tool that should be attractive for a wide range of applications.

## 2 Materials and Methods

### 2.1 Z-Splitter Prism

A schematic of our multifocus imaging microscope is shown in Fig. 1(a), which in essence is a standard widefield microscope but where the sample is first imaged to a z-splitter prism and subsequently re-imaged to a single camera. The key component here is the z-splitter prism, which splits the detection path into multiple paths of increasing pathlength such that each image is conjugated to a different depth in the sample.

**Figure 1.**
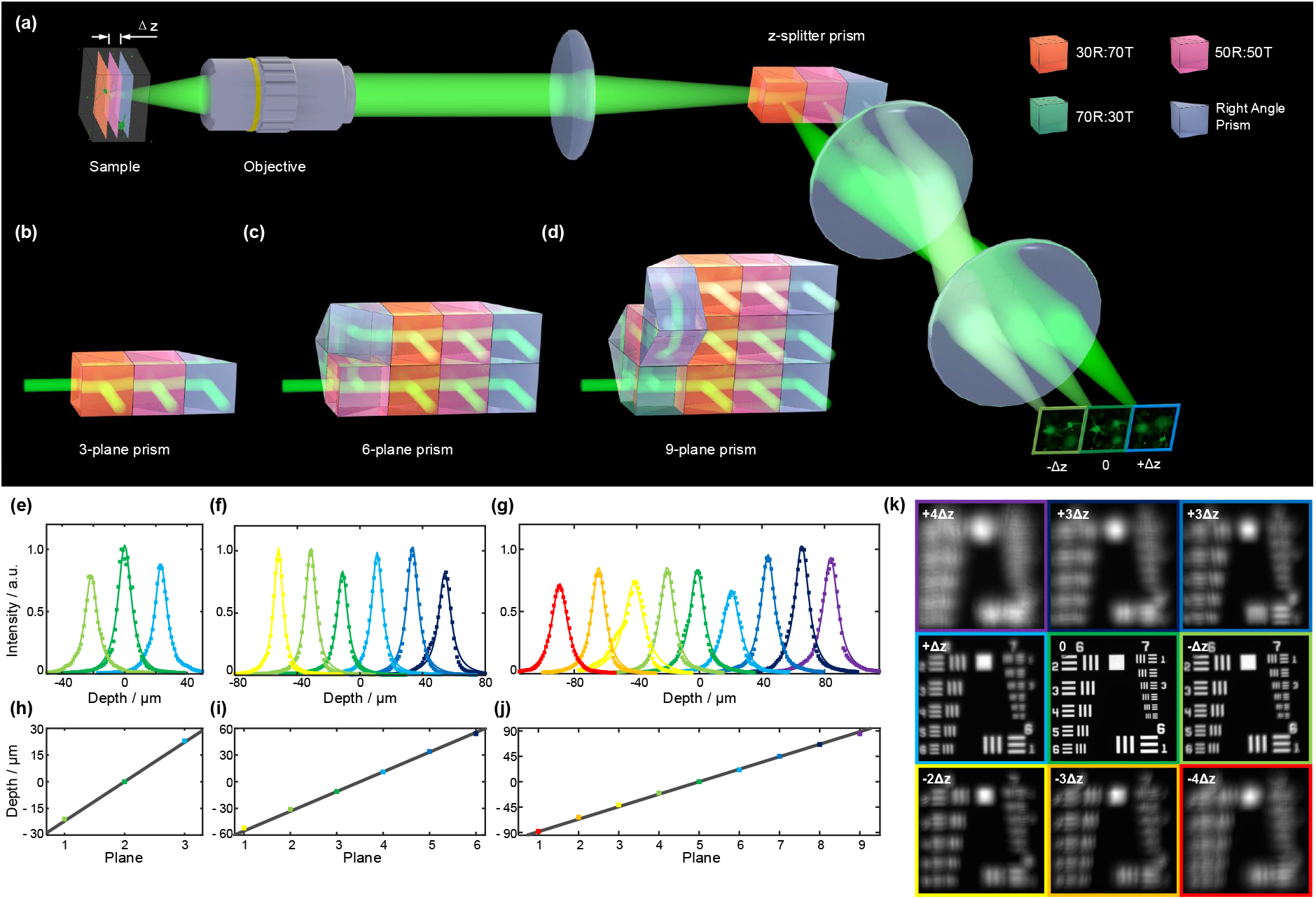
(a) Schematic of our multifocus imaging system. (b-d) Construction of 3-, 6- and 9-plane z-splitter prisms. (e-f) Fitted axial intensity profile of a 1 μm fluorescent bead measured with 3-, 6- and 9-plane prisms. (h-j) Theoretical and measured axial focal positions associated with 3-, 6- and 9-plane prisms. (k) Simultaneous 9-depth transmission imaging of a USAF resolution target using a 9-plane prism. Δ*z* = 22.1 μm.

The 3-plane prism (Fig. 1(b)) is the basis of all our z-splitter prisms. This consists of a 30R:70T beamsplitter (BS) cube, a 50R:50T BS cube and a right-angle prism all cemented together using optical adhesive. These beam splitting ratios are chosen so that roughly 1/3 of the light power exits from the output surface of each BS/prism. The optical pathlength associated with each output beam thus increases by *n_glass_L* upon transit through each successive BS, where *n_glass_* is the refractive index of the BS/prism glass and L is the length of each BS cube (assumed the same). The output beams are spatially distributed in a 1 ×3 geometry and directed onto a single camera through a telecentric 4*f* relay. By simple geometric optics, each focal plane in the sample is separated by a distance

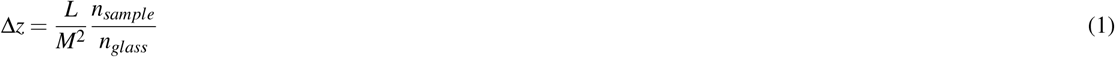

where *M* is the magnification from the sample to the z-splitter, and *n_sample_* is the refractive index of the sample. Similarly, the lateral FOV is given by

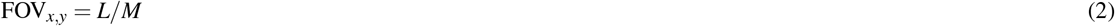

Thus by controlling the BS/prism dimension *L* and microscope magnification *M*, the lateral FOV and axial separation can be adjusted to the application of interest. Note that actual FOV can be somewhat smaller because the clear aperture of each beamsplitter can be smaller than *L*.

Similarly, we can construct a 6-plane prism (Fig. 1(c)) by splitting the incident beam into two paths using a single 50R:50T BS, with one beam optical pathlength extended an additional 3*n*_*glass*_*L* with respect to the other. The two 3-plane prisms arranged in two rows direct each path onto different regions of the camera, forming a 2 × 3 image of 6 focal planes. For the 9-plane prism (Fig. 1(d)), we use three 3-plane prisms that provide a 3 × 3 image geometry of 9 focal planes at the camera. Similarly, we can use a 70R:30T and a 50R:50T cube BS to split the path into three rows before entering the three 3-plane prisms, the optical pathlengths of the middle and top rows being extended respectively by 3*n*_glass_*L* and 6*n*_*glass*_*L* compared to the bottom row.

We assembled 3-, 6- and 9-plane z-splitter prisms with dimension *L* = 12.5 mm. Using a 20 × /0.5NA water immersion objective (*M* = 22.2), this leads to focal planes in the sample of axial separation Δ*z* = 22.1 μm and lateral span FOV_*x,y*_ = 1.25 mm. We measured the axial intensity profile of a 1 μm fluorescent bead (Fig. 1(e-g)), and plotted its optimal focus positions, which agree with theory (Fig. 1(h-j)). An example of 9-plane transmission imaging of a USAF resolution target is shown in Fig. 1(k).

An important feature of modern sCMOS cameras is their line-by-line readout structure, which enables increased frame rate by vertically cropping the sensor area of interest (AOI). Our z-splitter designs can take full advantage of this feature: while a 9-plane prism provides an overall square image geometry that is able to fit the full sensor area and run at full frame rate, a 3-plane prism, provides an elongated 1 : 3 imaging format, enabling higher imaging speed by vertical cropping to 1/3 of the sensor area. For a 6-plane prism, one can operate either at 1.5 times the full frame rate by using 2/3 of the sensor, or at 3 times the full frame rate by sacrificing half the vertical FOV and only using 1/3 of the sensor (see Fig. 5(c,d)). All these configurations can be swapped on site depending on whether one prefers a higher imaging speed or larger imaging volume.

### 2.2 Contrast Enhancement with Extended-Volume 3D Deconvolution

Because our multifocus fluorescence microscope is fundamentally based on standard widefield microscopy, it does not inherently reject out-of-focus background. A common strategy to mitigate this problem is to suppress out-of-focus background *a posteriori* by image deconvolution^33^, which seeks to reconstruct the (background-free) object distribution 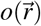 that satisfies

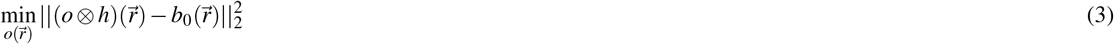

where 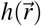 is the 3D PSF of the system, 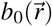 is the recorded 3D image stack, ⊗ denotes convolution, and 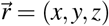 is the 3D Cartesian coordinate. It is important to note that the domain of both the image 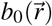 and estimated object 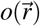 is confined to a cube defined by the imaging volume: 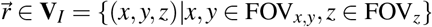, where FOV_*x,y*_ and FOV_*z*_ are the lateral and axial ranges of the imaging volume. This leads to the assumption that all signals observed in 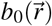 originate from sources within the imaging volume **V**_I_. However, this assumption is often invalid when the sample is thicker than the imaging volume and signals can originate from outside **V**_*I*_, i.e., far-out-of-focus fluorescence. Strategies have been reported to account for such contributions, such as assuming constant background^34^ or using 3D deconvolution applied to 2D images^35,36^.

Here, we present an extended-volume 3D deconvolution algorithm that takes full advantage of our captured 3D focal stack while also accounting for the possibility of additional out-of-focus fluorescence contributions from beyond the imaging volume. Our model considers the optimization problem

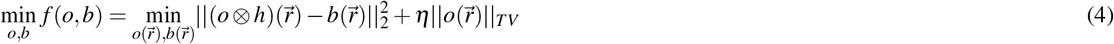

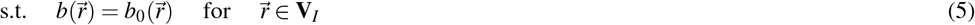

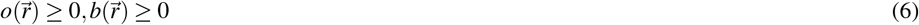

where 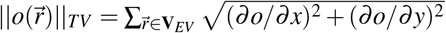 is the isotropic total variation norm in the lateral direction, and ***η*** is a small regularization parameter. The crux of our model is that we make use of a domain for *b, o, h* that is now 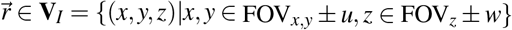, which is an expansion of **V**_*I*_ in all three spatial dimensions, where *u* ≥ 0 and *w* ≥ 0 denote the amounts of expansion in the lateral and axial directions. This simple expansion of the deconvolution domain allows us to assign signals in 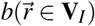 to sources in 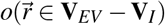, therefore enabling a better estimate of out-of-focus fluorescence. However, here both the object 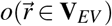 and the image 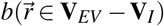 are unknown, so we adopt the block coordinate descent strategy, which alternately updates *o* and *b* through iterations. A schematic of this strategy applied to EV-3D is illustrated in Fig. 2(a), with the algorithm shown in Alg. 1.

**Figure 2.**
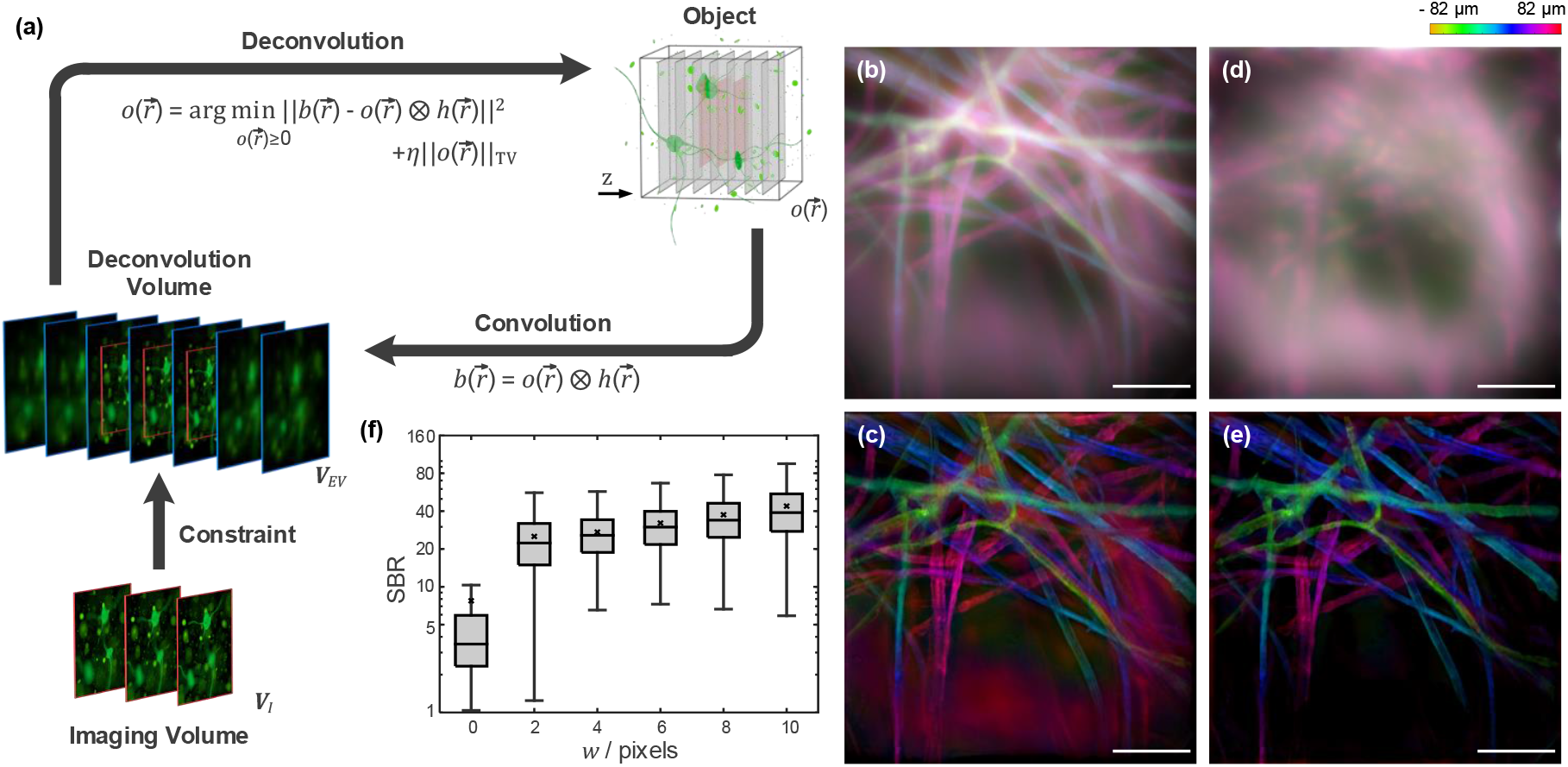
(a) Schematic of EV-3D deconvolution algorithm. (b) Color-coded EDOF image of the raw 9-plane image stack. (c) Color-coded EDOF image of the deconvolved stack using RL-3D algorithm. (d) EV-3D estimated background far-out-of-focus fluorescence in panel (b) generated from volume extension *V_EV_* – *V_I_*. (e) Color-coded EDOF image of the deconvolved stack using EV-3D algorithm. Scale bars are 100 μm. (f) SBR distribution as a function of axial extension w. Box plots: box, 25th to 75th percentiles; whiskers, 1.5× interquartile range from the 25th and 75th percentiles; middle horizontal line, median; ’×’ symbol, average.

Line 3 of Alg. 1 is essentially a deconvolution procedure that estimates *o* given image *b^k^* and *h*, the solution to which can be approximated by Richardson-Lucy deconvolution^37^. The nonnegativity constraint on 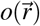 can be satisfied automatically with 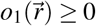, and *η* = 0. With the regularization parameter in the range 0 < *η* ≪ 1, small negative values might be introduced in 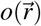. In such cases we set all negative values to 0. These steps are shown in Alg. 2, where *h*\* is the flipped PSF, and div(·) represents divergence.

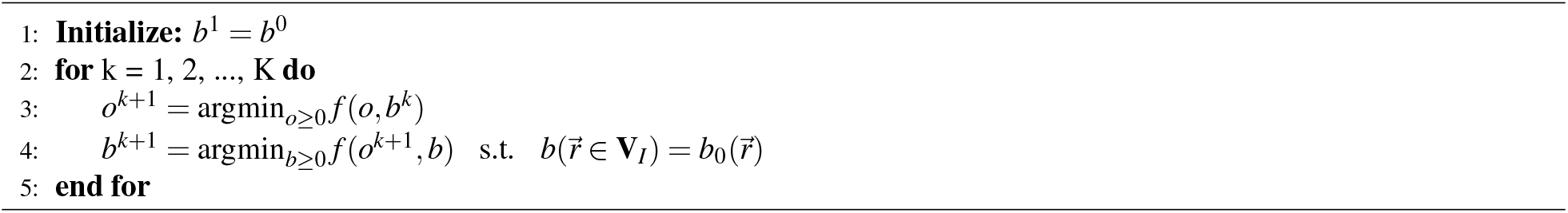

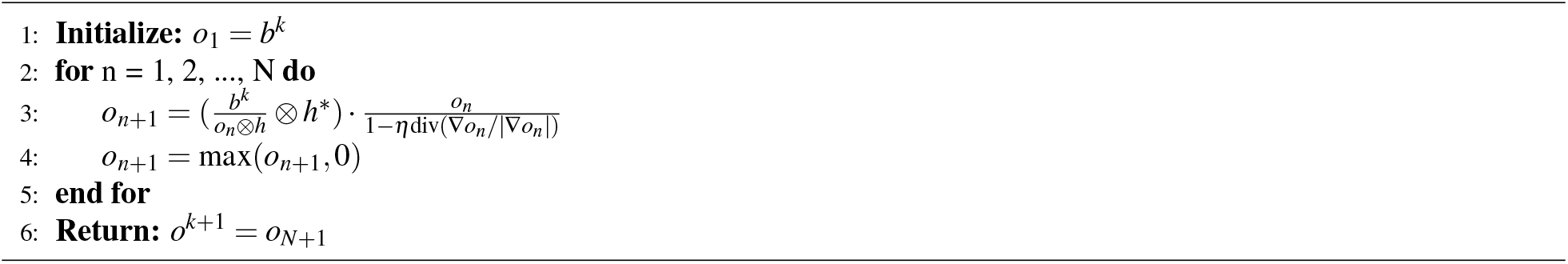

Line 4 in Alg. 1 is the forward convolution model, whose solution for step k can be explicitly expressed as:

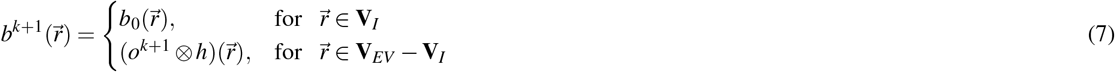

Note that this forward convolution operation also preserves nonnegativity.

As an example, we imaged a fluorescent lens-paper sample using a 9-plane z-splitter, obtaining a color-coded EDOF image shown in Fig. 2(b). Far out-of-focus background originating from outside the imaging volume is observable in the lower part of the image. Richardson-Lucy 3D (RL-3D) deconvolution, while reducing in-volume out-of-focus background, is unable to remove such far-out-of-focus background (Fig. 2(c)). However, with our new EV-3D algorithm, even far-out-of-focus background can be estimated, as shown in 2(d). With the removal of both in-volume and far-out-of-focus background, EV-3D results in higher reconstructed image contrast (Fig. 2(e)).

To quantitatively assess contrast improvement when using EV-3D deconvolution, we imaged a 1 mm thick high density fluorescent bead sample and measure the signal-to-background ratio (SBR) as a function of the deconvolution parameter w (Fig. 2(f)). Here, with a 20×/0.5NA objective and 6-plane z-splitter, the axial extent of *V_I_* was 110 μm, which was much smaller than the sample volume. The recorded image stack thus displays a large amount of out-of-volume background (see Fig. S3 for images before and after deconvolution). Without axial extension (*w* = 0), corresponding essentially to traditional RL-3D, the average SBR after deconvolution is 7.74, which is a substantial improvement over the raw image SBR of 0.75. An even better improvement comes from applying EV-3D deconvolution with an axial extension as small as 2 pixels (*w* = 2), in which case the average SBR jumps threefold to 25.1. Still further improvement is obtained with larger axial extension, though with diminishing returns: the average SBR is 27.2, 32.1, 37.4, 43.8 for *w* = 4,6,8,10, respectively. These results confirm the effectiveness of our EV-3D algorithm at improving image contrast.

A drawback of our EV-3D algorithm is its computation time, where each outer loop (iteration over k) requires an independent evaluation of RL deconvolution, which itself is iterative. A way to speed up the EV-3D algorithm is to employ a warm start strategy, where instead of initializing 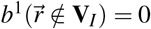, we initialize it somewhere close to optimal. One strategy, for example, is to initialize *b*^1^ as

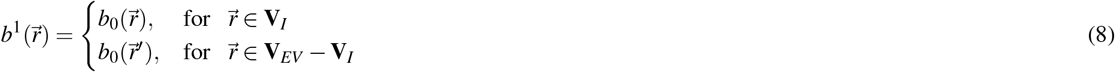

where 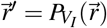 is the projection of 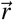 onto the set *V_I_*. This essentially assumes that 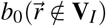 is equal to the value of the closest pixel within **V**_*I*_. Another strategy, applicable to video recordings with slowly varying background, is to initialize 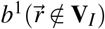 as the solution from the previous deconvolved frame. We have found that these warm-start strategies can increase the speed of EV-3D deconvolution more than an order of magnitude, while incurring only negligible degradation of reconstructed image quality (see Supplementary Fig. S2(e)).

For a typical deconvolution volume *V_EV_* of size 700 × 700 × 13 pixels, using outer iteration *K* = 20 and inner iteration *N* = 20, the algorithm requires 7.5 s per volume using Matlab 2019b on a desktop computer equipped with i9-9900K CPU, 64 GB RAM, and a RTX 2080 Ti GPU. Using warm start, the outer iteration is reduced to *K* = 1, and the net per volume computation time is thus reduced to 0.4 s.

## 3 Results

### 3.1 Multifocus Fluorescence Imaging

#### 3.1.1 Large-scale functional imaging in a mouse brain

The development of genetically encoded calcium^38^ and voltage^39^ indicators has enabled real-time monitoring of neural activity with a fluorescence microscope. However, brain function relies on the interactions of large populations of neurons distributed over extended volumes^40^. With a fluorescence microscope, while it is relatively easy to obtain a wide lateral FOV, signals away from the focal plane can be so blurred as to be irretrievable. However, with our multifocus imaging strategy combined with EV-3D deconvolution, we can easily record multiple focal planes across extended volumes with a single camera at high speeds, enabling high contrast, large scale functional imaging. We demonstrate this by performing *in vivo* calcium imaging in the mouse brain over volumes encompassing hundreds of neurons.

We imaged the striatum area of a mouse brain expressing GCaMP7 at a 50 Hz frame rate for 8 minutes (see Supplementary Video 1 for the complete recording after deconvolution). With a 3-plane z-splitter and a 10 × /0.28NA objective, we achieved a 3D FOV of 1.2 × 1.2 × 0.22 mm^3^ with an interplane separation of 110 μm. Note that while such an axial separation may appear high at first glance, we found it to be ideally suited for somatic imaging in the mouse brain. This is because the lack of inherent out-of-focus signal rejection, though generally considered a disadvantage in widefield microscopy, can actually be exploited here to image neuronal bodies beyond the microscope depth-of-field (defined by the Rayleigh range), albeit at reduced resolution. In our case, because somas in the striatum area are typically 10 μm in size^41^, they remain easily resolvable even when situated between focal planes.

After acquisition, we applied both RL-3D and EV-3D deconvolution to all frames for comparison, using theoretically calculated PSF. Figure 3(a) shows the temporal (max - min) projection of the 3 focal planes after deconvolution (see Supplementary Fig. S7 for a comparison of deconvolution results). We then made use of a custom-developed deep-learning-based segmentation procedure that identified 200, 345, and 186 neurons in each plane separately. Owing to the widefield nature of our microscope, some of these neurons appeared in more than one focal plane. The signals of such cases were merged, resulting in a total of 528 distinct neurons distributed throughout the imaging volume. The *ΔF/F* traces of neurons 250 - 349 during the 60 - 120 s recording period are shown in Fig. 3(b) (see Supplementary Fig. S8 for the complete 8 minute traces of all identified neurons).

**Figure 3.**
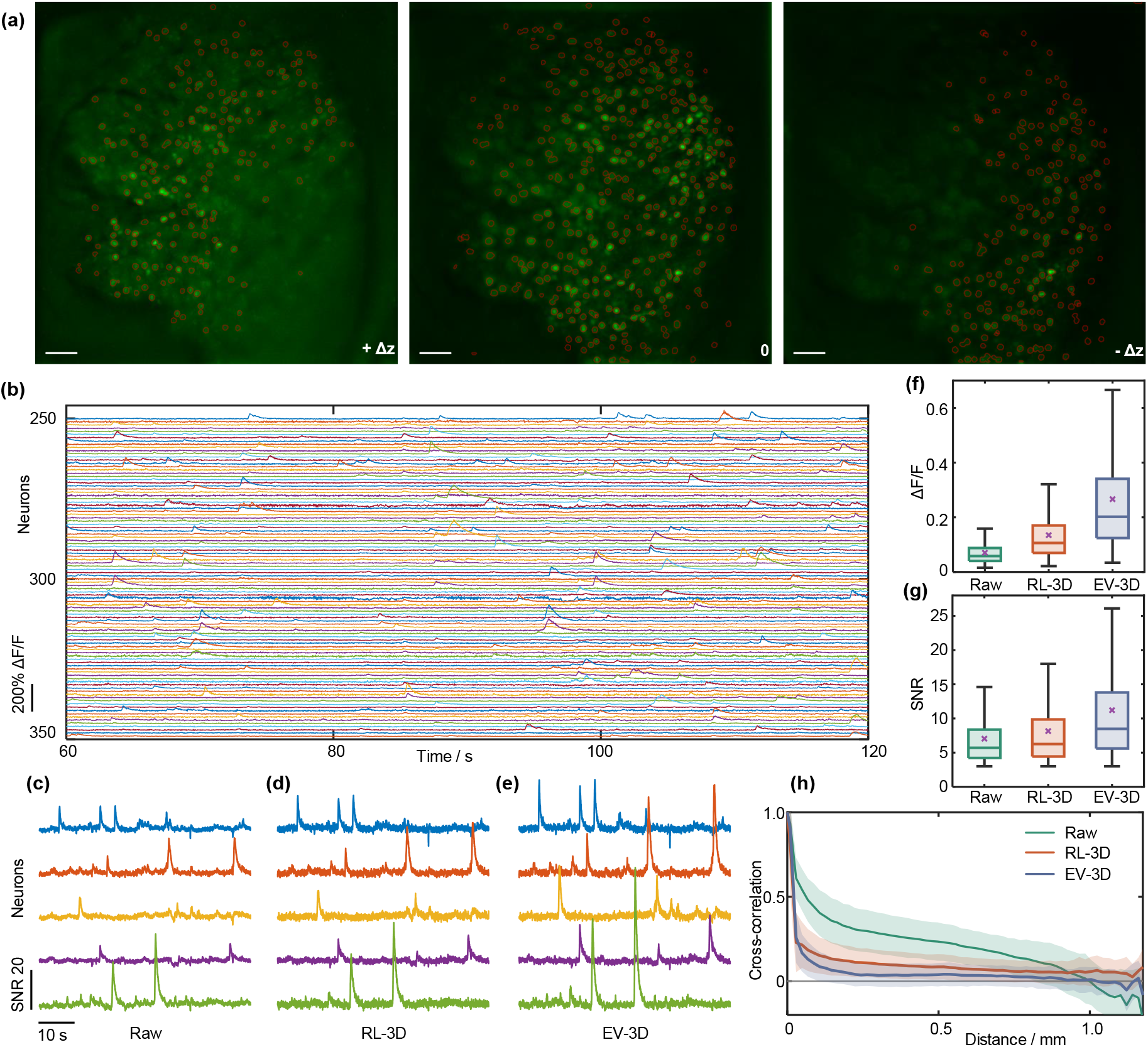
High-speed multifocus *in vivo* imaging of GCaMP7-labeled mouse brain over a large FOV. (a) (max - min) projections of three different focal planes over a 8 min recording at 50 Hz. The total 3D FOV is 1.2 × 1.2 × 0.22 mm^3^. From left to right, 200, 345 and 186 neurons are identified in each plane. Scale bar, 100 μm; Δ*z* = 110 μm. (b) Calcium traces of subset of neurons during 60 s - 120 s recording period. (c, d, e) Comparison of SNR for neurons 267 - 271 during 120 s −180 s recording period. (f) Distribution of Δ*F/F* conditioned on Δ*F/F* > 3*σ_n_* in cases of no deconvolution, RL-3D deconvolution and EV-3D deconvolution. (g) SNR distribution conditioned on Δ*F/F* > 3*σ_n_* in cases of no deconvolution, RL-3D deconvolution and EV-3D deconvolution. Box plots the same as Fig. 2(f). (h) Pairwise cross-correlation as a function of inter-neuron distance in cases of no deconvolution, RL-3D deconvolution and EV-3D deconvolution. Solid line, average cross-correlation; shaded area, ±1 std.

To quantify the improved performance of EV-3D compared to conventional RL-3D deconvolution, we evaluated both signal contrast (defined by Δ*F/F*) and SNR (defined by (Δ*F/F*)/*σ_n_*, where *σ_n_* is the noise standard deviation). In Fig. 3(c-e), we compared the SNR of 5 neuronal calcium traces over a 1 minute recording period extracted from the video. While the applications of RL-3D and EV-3D deconvolution both led to clearly improved image quality, the best improvement came from the latter, as manisfested by the increased SNR in calcium transients. This increase was consistent across all neurons within the FOV, where Figs. 3(f,g) encapsulate the statistics of common signal time points conditioned on Δ*F/F* > 3*σ_n_* without deconvolution, and with RL-3D and EV-3D deconvolution. On average, Δ*F/F* is increased from 0.070 to 0.135 with RL-3D, and is further increased to 0.266 with EV-3D. The average SNR is increased from 7.04 to 8.15 with RL-3D, and further increased to 11.2 with EV-3D.

A well-known problem associated with widefield imaging of large populations of neurons is signal cross-contamination, leading to more apparent neuronal signal correlations than actual^42^. Since out-of-focus light contributes to the majority of such cross-contamination, this problem is alleviated by the application of deconvolution. In Fig. 3(h) we show the pairwise cross-correlation of calcium traces as a function of inter-neuron separation distance before deconvolution, and after RL-3D and EV-3D deconvolution. We found that deconvolution reduced the pairwise cross-correlation towards 0, with the reduction being most significant for nearby neurons. On average the cross-correlation was reduced from 0.247 without deconvolution to 0.093 with RL-3D deconvolution, and further reduced to 0.043 with EV-3D deconvolution. This reduction was mostly consistent across all distances when comparing RL-3D and EV-3D results (orange and purple curves in Fig. 3(h)). Since actual neuron cross-correlation is typically close to zero^42^, these results suggest that a considerable amount of the observed cross-correlation in our images was erroneously induced by out-of-focus background, and that the application of EV-3D deconvolution was most effective at suppressing this background.

#### 3.1.2 Tracking freely moving Caenorhabditis Elegans

We further imaged cytoplasmic GCaMP6s signals in *C. elegan* neurons in a 510 × 510 × 170 μm^3^ 3D FOV with a 9-plane z-splitter and 20×/0.5NA objective. This enabled us to monitor, in 3D and at video-rate, multiple *C. elegans* undergoing free locomotion (Supplementary Video 2). A common method of *C. elegans* imaging makes use of spinning disk confocal microscopy^43,44^ to ensure both adequate imaging speed (video-rate) and high contrast (out-of-focus light rejection). Although our widefield multifocus strategy does not provide physical out-of-focus background rejection, it does, when complemented with EV-3D deconvolution, provide quasi optical-sectioning by numerical out-of-focus background suppression, enabling even fine details in the worms to be revealed with high contrast (Fig. 4(a-d)).

**Figure 4.**
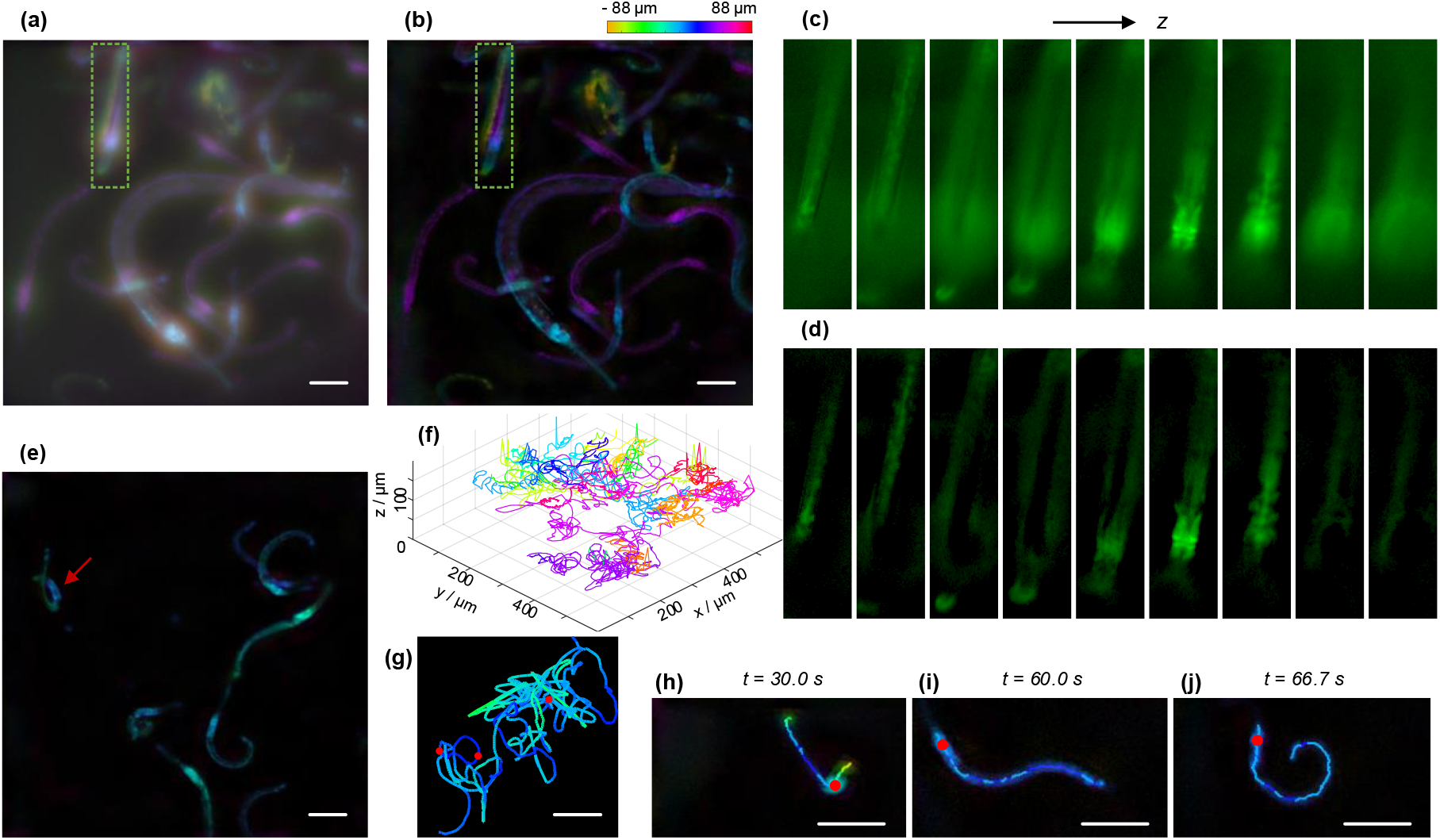
Video-rate multifocus tracking of freely moving *C. elegans.* (a) Color-coded EDOF image of one frame of the *C. elegans* video before deconvolution. (b) Color-coded EDOF image of the same frame after EV-3D deconvolution. (c,d) Individual 9-plane images over the green rectangular regions in (a) and (b) respectively. (e) One frame from the video used for *C. elegans* tracking. (f) All 27 traces of tracked head ganglia positions from a 104 s video. (g) Position of the head ganglia of a single worm (indicated by red arrow in (e)) from 12 s to 86 s. Line color represents depth. (h-j) Extracted skeletons of a single worm at times t = 30.0 s, 60.0 s and 66.7 s respectively. Red dots represent tracked head ganglia positions. Positions in (h-j) correspond to the rightmost, leftmost and middle red dotted positions in (g). All scale bars are 50 μm. For visualization, the gamma values of all images are set to 0.5.

An important application that is enabled by our system is 3D tracking throughout extended volumes. To demonstrate this, we recorded a 104 s video (Supplementary Video 3) at 30 Hz and tracked the head ganglia positions of worms over the entire duration of the video. Our large 3D FOV allowed multiple worms to be tracked at the same time, as shown in Fig. 4(f) where we plotted all 27 detected trajectories of head ganglia positions. Continuous tracking of a single worm is also possible, as shown in Fig. 4(g) where the trajectory of a single worm (indicated by the red arrow in Fig. 4(e)) is plotted over the course of 74 s. In addition to tracking, our system can be adapted to monitor whole body posture and locomotion-related behaviors in 3D^45^. In Fig. 4(h-j), we plotted the 3D body posture of a single worm at three different time points using skeleton analysis, revealing distinct crawling and swimming behaviors. These results were obtained at full frame rate and without the requirement of any mechanical scanning or active worm tracking^43,44^.

### 3.2 Multifocus Phase-Contrast Imaging

Our multifocus imaging strategy using a z-splitter is by no means limited to fluorescence imaging. For example, phase-contrast microscopy is widely used to visualize transparent biological samples^1^, and has the advantage of being label-free. A simple technique to obtain phase contrast is with the use of asymmetric illumination^46^. It can be shown that with such illumination, the 3D phase PSF inherently provides optical sectioning^47,48^. Thus, with multifocus imaging, 3D phase contrast can be obtained even without the aid of deconvolution.

To illustrate the versatility of our device we performed 3D phase-contrast imaging of living rotifers, a model organism for aging and ecology research^49^. Using a 9-plane z-splitter and 10 × /0.3NA objective, we imaged with a 1.1 × 1.1 × 0.7 mm^3^ 3D FOV at 30 Hz. This combination of FOV and imaging speed allowed us to observe the behavior of multiple rotifers in their natural state (Fig. 5(a) and Supplementary Video 4). High speed imaging over such a large 3D FOV would have been difficult with alternative multiplane imaging systems. We note that because the sample here was partially absorbing, our images contain both absorption and phase contrast. In principle, these contrasts can be isolated by sequential imaging with complementary illumination apertures^46,50^. We did not implement this strategy, however, because of the attendant reduction in speed. In general, a trade-off between 3D FOV and spatial resolution can be effected in our system by simply switching objectives. For example, by switching to a 40 × /0.8NA objective, we could readily resolve internal tissue structures such as the stomach, mastax and corona across different focal planes (Fig. 5(f-h)), albeit at a reduced 3D FOV of 280 × 280 × 44 μm^3^.

**Figure 5.**
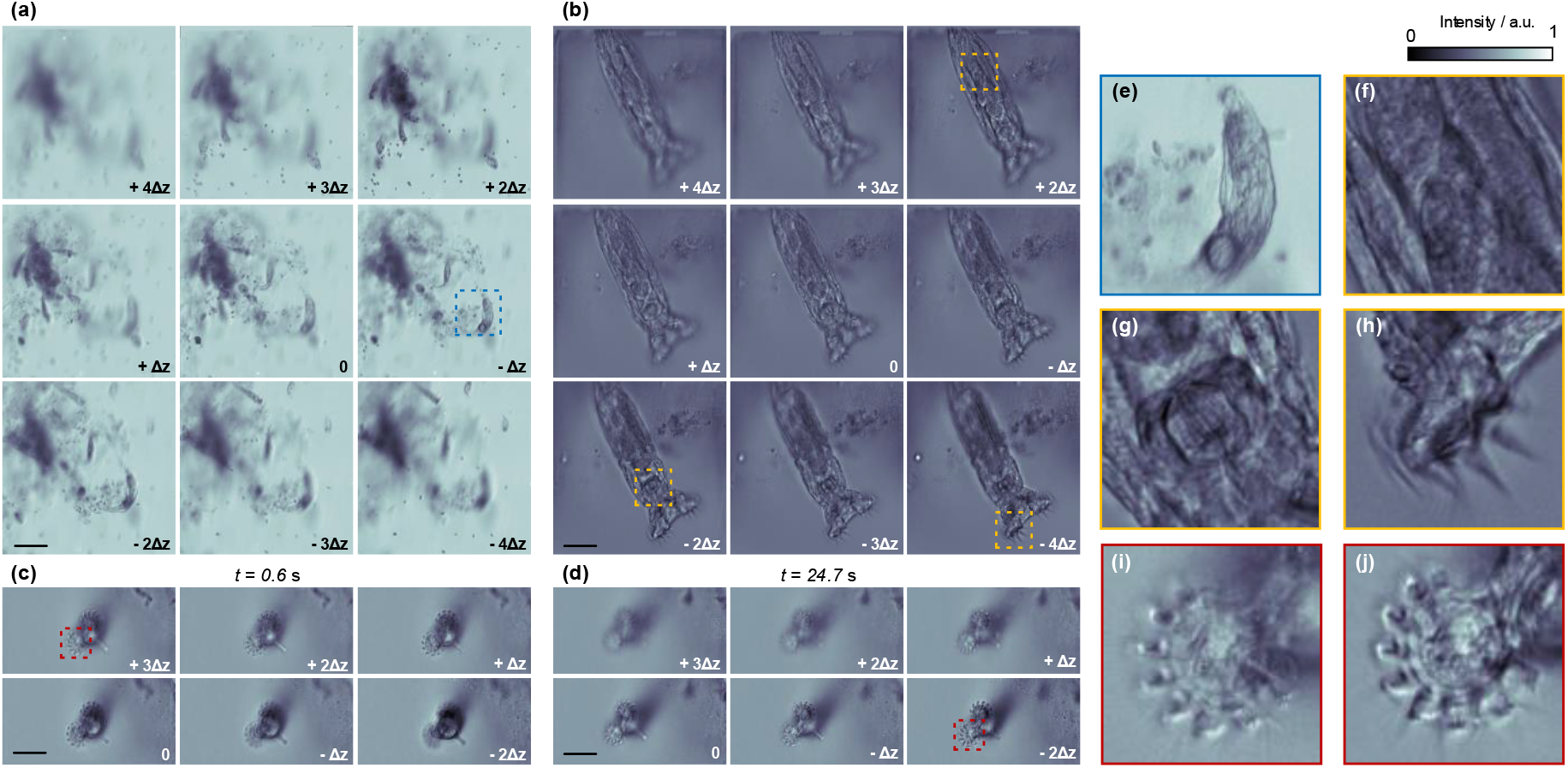
Multifocus phase-contrast imaging of living rotifers. (a) Large FOV (1.1 × 1.1 × 0.7 mm^3^) imaging of multiple rotifers in their natural state. Δ*z* = 88.6 μm. Scale bar, 200 μm. (b) Multifocus image of a single rotifer at high resolution. Δ*z* = 5.5 μm. Scale bar, 50 μm. (c,d) High speed imaging (100 Hz) of beating cilia of a rotifer. (c) and (d) show two frames at times *t* = 0.6 s and *t* = 24.7 s, respectively. Δ*z* = 5.5 μm. Scale bar, 50 μm. (e) Expanded view of a single rotifer within the blue square in (a). (f-h) Expanded view of stomach, mastax and corona regions (yellows squares at depths +2Δ*z*, −2Δ*z* and −4Δ*z* in (b)) of a rotifer at different depths. (i,j) Expanded view of cilia from the red square regions in (c) and (d), respectively. The focal position of the cilia moved from +3Δ*z* at *t* = 0.6 s to −2Δ*z* at *t* = 27.4 s. a.u., arbitrary unit.

In the above experiment, the acquisition speed was limited by the camera full frame rate, meaning we could observe entire living organisms at video-rate (Supplementary Video 5). However, even higher acquisition speed is desired when imaging the beating of cilia on the corona (with which rotifers create water currents to draw in food^49^). The ciliary beat frequency can be up to 30 Hz, prescribing high speed imaging to reveal any changes in frequency^51^. Previous techniques based on single-plane imaging impose strict limitations on sample movement, as well as coronal orientation, which must be parallel to the focal plane^51,52^. Here, by using a 6-plane z-splitter and cropping the camera vertical AOI, we imaged the cilia beating across different depths at 100 Hz over a 3D FOV of 280 × 140 × 28 μm^3^ without motion blur (Supplementary Video 6). Because the imaging volume was larger than the size of the corona, we were further able to accommodate global animal motions without active tracking: although the cilia swept from +3Δ*z* at *t* = 0.6 s to −2Δ*z* at *t* = 27.4 s (see Fig. 5(c,d) and expanded views in Fig. 5(i,j)), they could still be resolved by virtue of multifocal imaging.

### 3.3 Multifocus Darkfield Imaging

Darkfield microscopy is yet another modality routinely used to for high contrast imaging of unstained biological samples^1^. Adaptation of our system to 3D darkfield imaging is straightforward: one need only insert an annular ring behind the condenser to illuminate the sample from oblique angles larger than the detection numerical aperture.

With our system, volumetric samples can be imaged in a single shot. Figure 6(a-c) shows all-in-focus images of radiolaria, volvox, and dictydium obtained with a 9-plane z-splitter (the individual 9 plane images are shown in Supplementary Fig. S9). Such samples typically extend hundreds of microns in all dimensions, making them difficult to capture in a single frame with a standard darkfield microscope. However, with a 20 × /0.5NA objective we can achieve a 520 × 520 × 200 μm^3^ FOV, enabling the rendering of all-in-focus images that span the entire specimens.

**Figure 6.**
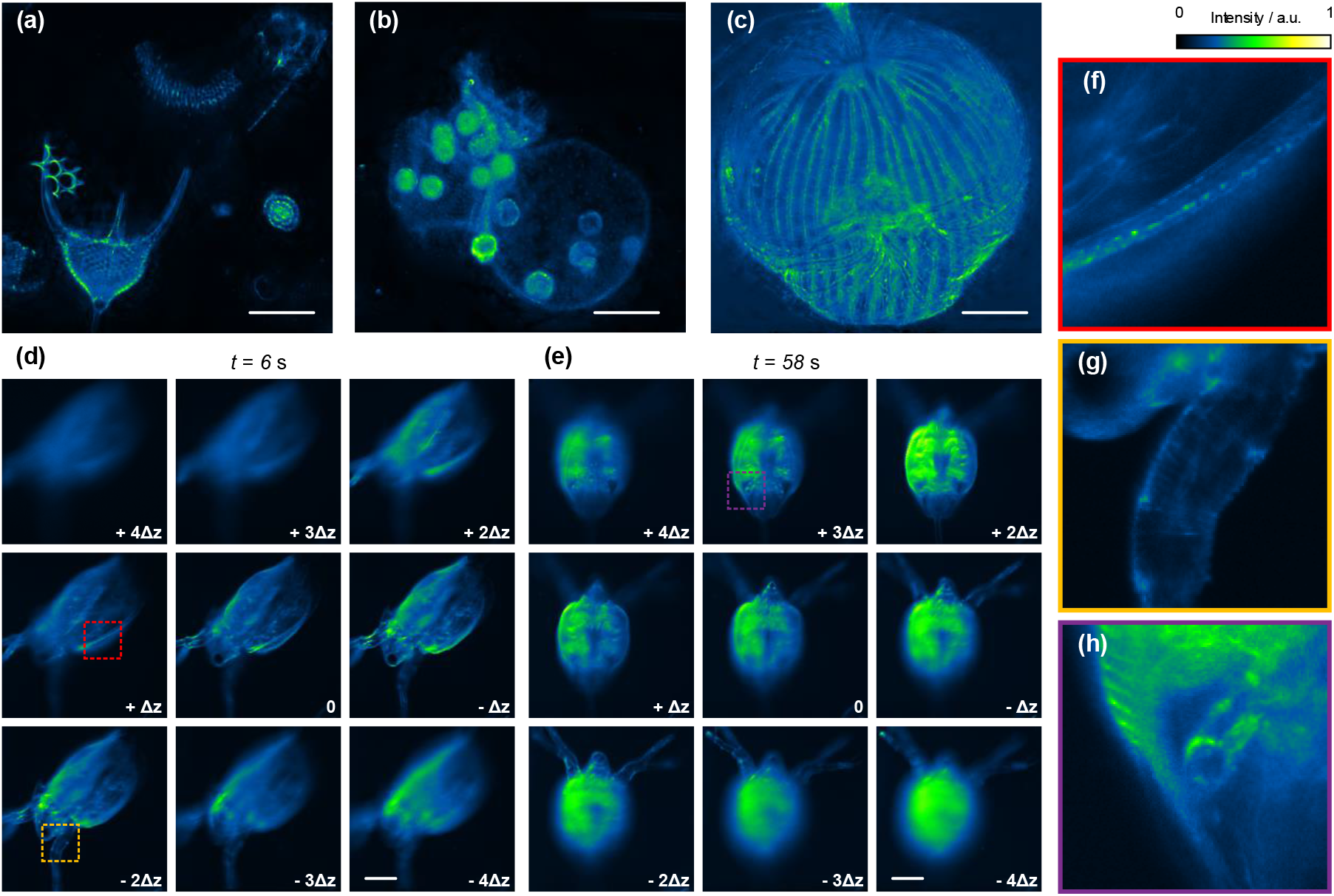
Multifocus darkfield imaging of entire organisms. (a - c) All-in-focus images of radiolaria, volvox, and dictydium. Scale bar, 100 μm. (d, e) 9-plane images of a living *Daphnia magna* at times *t* = 6 s and *t* = 58 s, respectively. Δ*z* = 88.6 μm. Scale bar, 200 μm. (f-h) Expanded views of boxes in (d, e), where distinct features can be seen across depths. a.u., arbitrary unit.

Similar to phase-contrast imaging, here we can also image an entire behaving animal at full frame rate. As a demonstration, we imaged a living *Daphnia magna* for 74 s at 30 Hz. By virtue of our large 3D FOV (1.1 × 1.1 × 0.7 mm^3^), the animal did not need to be immobilized before imaging. This allowed free movement of the animal within the imaging chamber, where, although the animal abruptly changed posture several times during the recording, we were still able to image its body in its entirety (Fig. 6(d,e)). The complete recording is provided in Supplementary Video 7.

## 4 Discussion

Our multifocus imaging strategy offers several notable advantages over alternative strategies. Compared to techniques using a DOE^29–31^, our z-splitter prism is low cost, easy to design and assemble, does not require the addition of a chromatic corrector, and can be adapted to a variety of imaging modalities. Compared to techniques that also involve the use of beamsplitting optics^24,25,27,28^, our system requires only a single camera and enables significantly larger FOV while offering the flexibility of swapping in different z-splitters to accommodate different imaging speed or volume requirements. For example, our imaging speed is solely limited by the camera frame rate. Elongated z-splitter geometries can take advantage of the readout structure of sCMOS cameras to achieve faster speeds by AOI cropping. We demonstrated 100 Hz imaging with a 2 × 3 z-splitter using a camera of only modest speed (30 Hz full frame rate). Higher speeds still could easily be attained with faster cameras. We also note that different z-splitter geometries can be considered, abiding by the same design philosophy. As an example, we provide a 12-plane design in Supplementary Document. Other designs such as 1 × 4 or 2 × 2 geometries are also possible with off-the-shelf components.

In addition to the hardware advantage that comes from the use of a z-splitter prism, we introduced a software advantage that comes from an extended-volume 3D deconvolution algorithm that specifically exploits the multiplane image information provided by our system. This algorithm allocates the reconstructed distribution of fluorescent sources into an extrapolated volume, thus making allowances for far-out-of-focus background and leading to significantly improved contrast and SNR of our widefield fluorescence images compared not only to the raw multifocus images but also to a more conventional 3D deconvolution algorithm that distributes the sources only in a confined volume. We anticipate that this algorithm will be greatly beneficial in imaging applications involving thick samples where the fluorescence labeling is dense, as is often encountered, for example, when imaging brain tissue.

Finally, owing in a large part to the geometric simplicity and achromatic nature of our z-splitter prism, we show that our multifocus technique can be generalized to imaging modalities that go beyond fluorescence. Here we demonstrated the additional possibilities of fast, 3D phase-contrast and darkfield imaging over large scales, but still other modalities can be considered. For example, with regard to 3D phase imaging, currently we employ only a single oblique illumination angle to obtain 3D phase-gradient contrast. However, *quantitative* 3D phase imaging is possible by combining multiple oblique illumination angles with deconvolution^46,50^, possibly aided by strategies invoking the transport of intensity equation^53^ or iterative algorithms^54^. Simple modifications to the illumination or detection paths could also enable possibilities of spectral^25,29^ or polarization^30^ imaging in 3D.

In summary, we have demonstrated a flexible and versatile platform for high speed. high contrast, large field-of-view 3D imaging that makes use of a simple z-splitter implemented with standard widefield microscopy. The platform can be applied to a diverse range of imaging applications, making it an attractive tool for biological and biomedical research.

## Acknowledgments

The authors thank G. Wirak and C. Gabel for help in C. elegans culturing.

## Additional Information

See Supplementary Document for supporting content.

